# Structural Basis for Hyperpolarization-dependent Opening of the Human HCN1 Channel

**DOI:** 10.1101/2023.08.17.553623

**Authors:** Verena Burtscher, Jonathan Mount, John Cowgill, Yongchang Chang, Kathleen Bickel, Peng Yuan, Baron Chanda

**Affiliations:** Department of Anesthesiology, Washington University School of Medicine, Saint Louis, MO, USA; Department of Cell Biology and Physiology, Washington University School of Medicine, Saint Louis, MO, USA; Center for the Investigation of Membrane Excitability Diseases, Washington University School of Medicine, Saint Louis, MO, USA; Department of Biochemistry and Molecular Biophysics, Washington University School of Medicine, Saint Louis, MO, USA; Department of Neuroscience, Washington University School of Medicine, Saint Louis, MO, USA; Department of Applied Physics, KTH Royal Institute of Technology, Stockholm, Sweden; Department of Pharmacological Sciences, Icahn School of Medicine at Mount Sinai, New York, NY, USA; Department of Neuroscience, Icahn School of Medicine at Mount Sinai, New York, NY, USA

## Abstract

Hyperpolarization and cyclic-nucleotide (HCN) activated ion channels play a critical role in generating self-propagating action potentials in pacemaking and rhythmic electrical circuits in the human body. Unlike most voltage-gated ion channels, the HCN channels activate upon membrane hyperpolarization, but the structural mechanisms underlying this gating behavior remain unclear. Here, we present cryo-electron microscopy structures of human HCN1 in Closed, Intermediate, and Open states. Our structures reveal that the inward motion of two gating charges past the charge transfer center (CTC) and concomitant tilting of the S5 helix drives the opening of the central pore. In the intermediate state structure, a single gating charge is positioned below the CTC and the pore appears closed, whereas in the open state structure, both charges move past CTC and the pore is fully open. Remarkably, the downward motion of the voltage sensor is accompanied by progressive unwinding of the inner end of S4 and S5 helices disrupting the tight gating interface that stabilizes the Closed state structure. This “melting” transition at the intracellular gating interface leads to a concerted iris-like displacement of S5 and S6 helices, resulting in pore opening. These findings reveal key structural features that are likely to underlie reversed voltage-dependence of HCN channels.

## INTRODUCTION

The voltage-dependent ion channels with cyclic nucleotide binding domains (CNBD) on their C-terminal end are a family of voltage-gated ion channels (VGICs) that exhibit unusual diversity in their voltage-dependent activity. These channels respond to depolarizing membrane voltages ^1,2^, but some are virtually insensitive^3–6^ or respond to hyperpolarizing membrane voltages^7,8^. The hyperpolarization and cyclic nucleotide gated (HCN) ion channels are a class of ion channels within the CNBD family that open when membrane voltage is more negative than a typical resting membrane potential. This unique property is crucial for their physiological role in pacemaking and spike synchronization^9^ in the heart and nervous system^8,10,11^. HCN channels are also regulated by cyclic nucleotides, which act via the CNBD, thereby fine-tuning the frequency of their pacemaking activity. As a result, HCN channels play a crucial role in integrating electrical signaling with a key second messenger pathway^12^.

The CNBD family of ion channels shares a similar tetrameric architecture with other members of the VGIC superfamily^13^. The first four transmembrane segments constitute the voltage-sensing domain (VSD) where the fourth helix (S4) contains a repeating pattern of positive charges at every third position. These charges are the primary sensors of membrane potential. The S5 and S6 helices of each subunit come together to form the central ion-conducting domain. The cytosolic CNBD domain in each subunit is connected to the S6 helix by the C-linker region. Unlike canonical VGICs, the CNBD family exhibits a non-domain swapped arrangement wherein VSD of each subunit is located next to its own pore helices^14–17^. These channels lack the extended S4-S5 linker helix that is critical for electromechanical coupling in domain-swapped ion channels. Comparison of resting and activated state structures of these channels reveals a mechanism of electromechanical coupling for depolarization-activation ion channels. The downward movement of the voltage-sensing S4 helix, upon hyperpolarization, pushes the S4-S5 linker towards the intracellular side. The S4-S5 linker acting as a lever arm holds the pore gates closed at negative voltages by exerting force on the adjacent S5 helix, which is relieved when the voltage sensor returns to Up position at depolarized potentials.

Electromechanical (EM) coupling in non-domain swapped ion channels must involve an alternate pathway given the absence of a S4-S5 linker helix in these channels. This non-canonical pathway of channel activation is also remarkably versatile, compared to domain swapped channels, which are all outwardly rectifying. Structures of depolarization activated EAG channels with voltage sensors in Down or Up configuration show that the pore is closed in both cases^14,18^. The central pore is also occluded in the structures of HCN channels where the voltage sensors are either in Up or Down conformation^17,19^. The S4 helix, in both channels, exhibits a break in the middle in the Down conformation suggesting that this is a distinct mechanistic feature of HCN channels^19,20^. Nevertheless, largely due to a lack of structures corresponding to hyperpolarization activated Open state, the central question as to how the HCN voltage-sensor in the Down state drives pore-opening remains unclear. A recent structure of HCN4 isoform with an Open pore and resting voltage sensors likely corresponds to a rare intermediate state rather than a bona fide open HCN channel^21^. In this study, we report three structures of a human HCN1 isoform in Open, Intermediate, and Closed conformations, revealing how progressive changes in the S4 and S5 helices result in pore opening and providing critical new insight into the mechanism of reversed gating polarity in HCN channels.

## RESULTS

To determine the structures of hyperpolarization-activated ion channels in various gating states, we focused our investigation on a human HCN1-F186C-S264C channel mutant, which was first described by MacKinnon and colleagues^19^. Metal cross-bridging of the two introduced cysteines clearly locks the voltage-sensing domain in an activated conformation as the S4 helix is shifted down by two helical turns compared with the position of S4 in the resting state. Remarkably, the S4 helix in this structure is interrupted by a break in the middle causing the lower part of this helix to become almost parallel to the membrane plane. Although this mutant is functionally locked open by metal cross-bridging in biological membranes, the pore helices are in a closed conformation in the cryo-EM structure determined in the presence of heavy metal. Thus, this structure corresponds to a putative intermediate or a pre-open state (PDB: 6UQF) where the voltage sensor is activated but the pore is closed.

Saponaro and colleagues found that the choice of solubilizing detergent drastically influences the conformation of the pore domain for the rabbit HCN4 isoform^21^. They speculated that DDM stabilizes the closed state whereas a mixture of LMNG and cholesterol hemisuccinate (CHS) renders the pore domain open even in the absence of voltage-sensor activation or ligand binding. Similarly, we wondered whether the detergents might also influence the pre-open state structure and provide sufficient stabilization to drive channel opening. To this end, we purified the HCN1-F186C-S264C mutant in a mixture of 0.025% LMNG, soy polar lipids and cholesterol hemi-succinate and determined its structure in the presence of Hg^2+^ to a global resolution of 3.73 Å using single-particle cryo-EM (Fig. 1a, right panel, Extended Data Figs. 1 and 2, Extended Data Table 1 and Movie 1). In the cryo-EM reconstruction, the cytosolic region, including the CNBD, is well resolved with clear side-chain densities for the majority of amino acids. However, while transmembrane helices S5 and S6 are well-resolved, the other four-especially S1 to S3- are at lower resolutions. Nonetheless, the unique, bent geometries of S1-S2 are consistent with the closed state structure and consistent with rigid-body motion, while densities for bulky side chains such as R270 and W281 along with the position of the cysteine metal bridge facilitates unambiguous registry (Extended Data Fig. 2). S3 appears to have undergone a conformational change in the Open state, and its corresponding density is better fit by a continuous helix than the S-shaped geometry observed in the closed structure. The activated voltage-sensor HCN model (PDB: 6UQF) was used as the initial model to build the atomic model of the open state HCN1 channel.

**Fig. 1:**
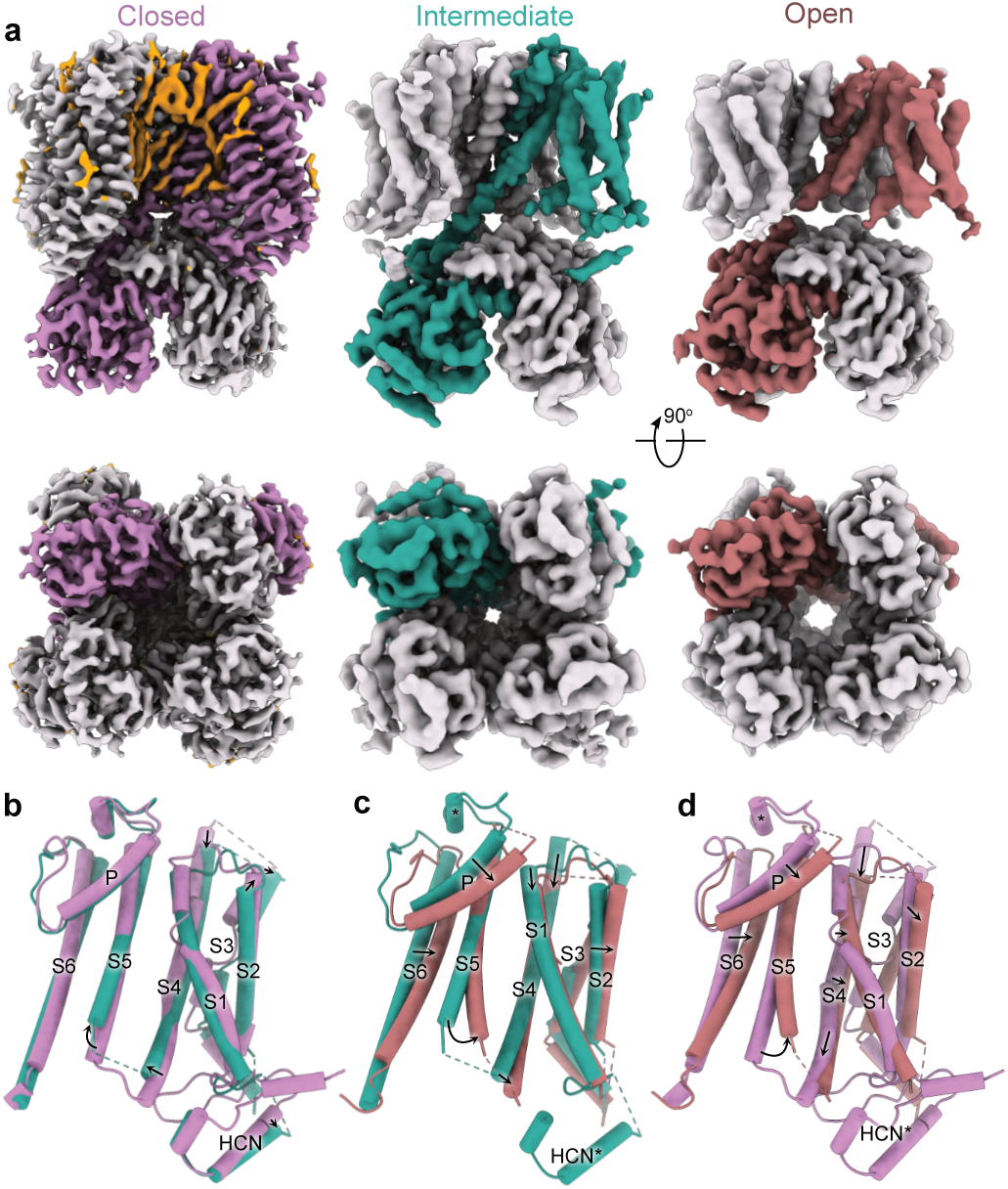
Structures of hHCN1 in Closed, Intermediate, and Open conformations. **a**, Cryo-EM density maps of corresponding conformations in side view (top panels) and bottom view (bottom panels). One protomer of each conformational state is colored either lilac (Closed), turquoise (Intermediate) or burnt red (Open). Putative lipid densities were only observed in the Closed state and are colored orange. **b-d**, Superposition of the transmembrane region corresponding to protomers in various conformations: the Closed and the Intermediate state (**b**); the Intermediate and the Open state (**c**); and the Closed and Open state (**d**).

Comparison with the structure of HCN4 with an open pore, in particular the pore-lining S6 helices, indicates that our HCN1-F186C-S264C structure represents an open conformation (Fig. 1a, Extended Data Fig. 3). Examination of the bottom view of the cryo-EM density map shows that the pore is dilated and the pore-lining S6 helices are displaced outward away from the central pore axis. The minimum distance of two opposing S6 helices, measured between the Cα atoms of the inner gate forming residue Q398, increased from 13.8 to 17.1 Å.

To ascertain the role of detergent-lipid mixture as opposed to metal cross-bridge in stabilizing the open conformation, we also solved the structure of this mutant under reducing conditions without changing the detergent-lipid mixture to a resolution of 3.26 Å (Fig. 1a, left panel, Extended Data Figs. 4 and 5, Extended Data Table 1 and Movie 2). This structure is nearly identical to the closed state structure reported previously (PDB: 5U6P)^17^. The voltage-sensing S4 helix is in Up position and the central ion conduction pathway is closed. Thus, we conclude that the Open channel conformation is primarily facilitated by oxidizing conditions that trap the voltage sensors in a hyperpolarized conformation.

Comparison of the Open and Closed structures reveals that while the open channel conformation involves tilting of S2, straightening of S3, and downward translocation of S1 and S4, the S1-S4 bundle itself is not radially displaced (Fig. 1d). Unlike in the bacterial cyclic nucleotide-gated ion channel, there is no significant upward movement of the CNBD domains towards the transmembrane portion^22^. We observed a number of lipid densities in the closed state that were absent in the open state, partly due to the lower resolution of the open state structure. Nevertheless, superposition of the Open and Closed state structures reveals that the movement of the pore helix in the open state would cause the displacement of the annular lipids observed in closed state structure (Extended Data Fig. 6), indicating that opening must be accompanied by rearrangement of the surrounding lipids.

Upon further examination of the cryo-EM densities, we discovered that C309 in the S5 helix and C385 in S6 likely form a disulfide bridge in the Open state but not in the Closed (Extended Data Figs. 2 and 5). Both cysteines are conserved in mammalian HCN isoforms, and substituting C309 with alanine results in a functional channel with a voltage dependence that is shifted towards more negative potentials indicating that it is more difficult to open the mutant channels (Extended Data Fig. 7). To obtain a more complete estimate of the energetic cost of disulfide crossbridge in stabilizing the Open state, we will need to measure open channel probability (Po), which is technically challenging given the low single-channel conductance^23^ of these channels.

To investigate the role of these cysteines further, we solved the structure of the HCN1-F186C-S264C-C309A mutant in the presence of Hg^2+^ and an identical LMNG:lipid:CHS ratio to a global resolution of 3.89 Å (Fig. 1a, middle panel, Extended Data Figs. 8 and 9, Extended Data Table 1 and Movie 3). Despite lower global resolution, the map is of higher overall quality than the Open structure and has fewer regions with broken density. Based on the position of the S4 and S5 helices, we classify this as the Intermediate state structure (Fig. 1a and b). Interestingly, we observe far fewer lipid densities in this intermediate state structure, with only 1-2 prominent densities nestled at the S5-S6 interface. Finally, it is worth noting that in both structures, we observe only weak densities corresponding to the N-terminal HCN domain, indicating that interactions with this domain and the CNBD are weakened upon activation.

Although the voltage-dependent activity of HCN channels is reversed, the voltage-sensing S4 segment moves in the same direction as depolarization-activated ion channels^19^. In other words, membrane hyperpolarization drives downward movements of S4 segments in all voltage-gated ion channels. A comparison of the voltage-sensing domains in the three HCN channel structures reveals that in the Intermediate state, the R4 charge of the S4 helix moves past the Cα position of the charge transfer center F186 (Fig. 2a). In the Open state, the R3 charge also moves past this position (Fig. 2b). The position of the voltage-sensing charges in the Open state is similar to that observed in the activated voltage-sensor structure corresponding to the pre-open state (PDB: 6UQF)^19^. The S4 helix retains its mid-helical bend in the Open state. However, unlike the activated state pre-open structure, it does not break or become parallel to the membrane plane (Fig. 2c). The degree of bend is slightly more pronounced than in the Down state structure of the EAG1 obtained in hyperpolarized membranes (Fig. 2d; PDB: 8EP1)^18^. Another unique feature is that the S4 helix is shorter by about two helical turns in the Open state structures, potentially due to unwinding of the intracellular end. We also see that the S4 helix shortens in the Intermediate state, but to a lesser extent than in the Open state structure. This helix unwinding is not observed in the activated voltage-sensor structure – despite being positioned similarly – or in the Down state structure of EAG1^18^, but was recently seen in the Down state structure of Nav 1.7^24^. Despite the shortening of the HCN1 S4 helix in the Down state, it still extends at least two helical turns further than the EAG1 S4 helix in the down conformation.

**Fig. 2:**
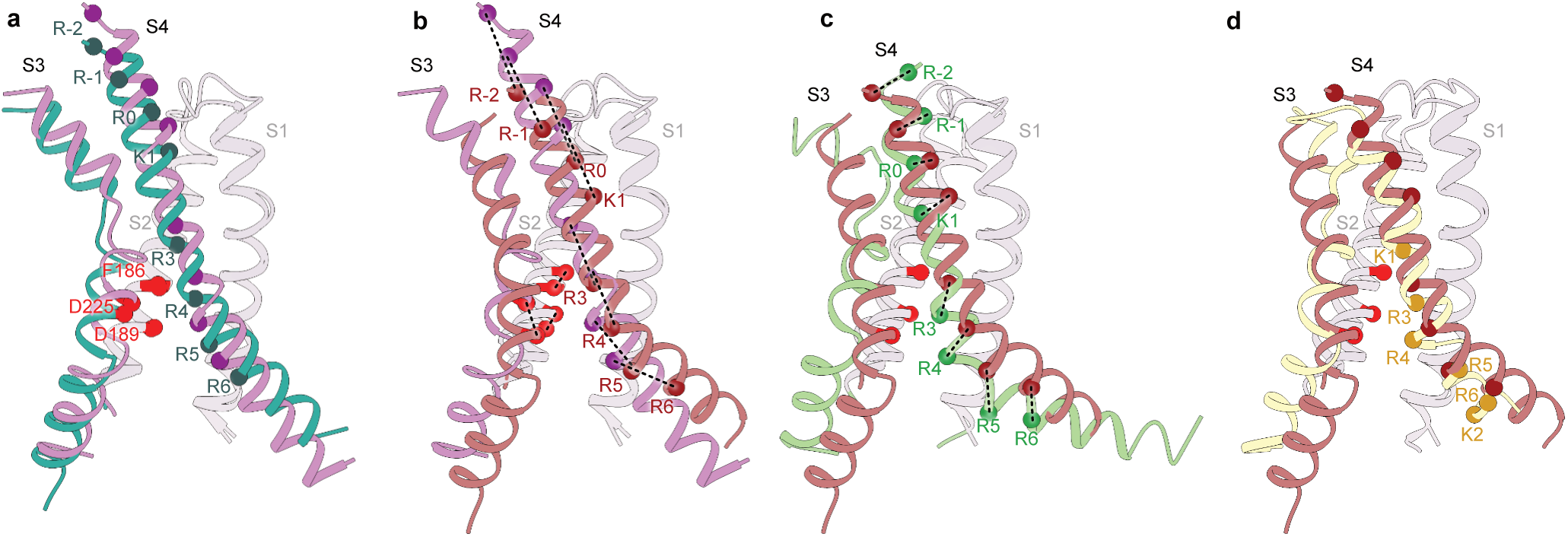
Movements of the S4 helix in the voltage sensing domains (S1-S4). **a-b**, Comparison of the S4 helix position between **a**, the Closed and Intermediate state and **b**, the Closed and Open state. The Cα atoms of positively charged residues in the S4 helix are depicted as spheres and are labelled as R-2 (R252), R-1 (R255), R0 (R258), K1 (K261), R3 (R267), R4 (R270), R5 (R273), R6 (R276). The Cα atoms of the residues F186, D225 and D189 constituting the charge transfer center are represented by red spheres. Dashed lines indicate the displacement of each charge. **c-d**, Comparison of the S4 helix position between the HCN1 Open state with either (**c**) the voltage sensor activated pore structure of HCN1 (6UQF) or (**d**) the closed pore EAG1 structure with the voltage sensor in the resting state (8EP1). Cα atoms of positively charged residues in the S4 helix of the open EAG1 channel are depicted as spheres and labeled as K1 (K327), R3 (R330), R4 (R333), R5 (R336), R6 (R339), and K2 (K340). All the structures were aligned with respect to the S1 and S2 helices.

To investigate the changes in structure related to electromechanical coupling, we focused on the S4 and S5 helices at the gating interface of all three structures (Fig. 3). In the Closed state of all hyperpolarization-activated ion channels, the S4 and S5 helices are close together and connected by a short loop^19,21,25^. Mutations in this crucial gating interface have been shown to disrupt the ability of the channels to close^26^. Strikingly, in the Intermediate and Open state structures, the intracellular helix-turn-helix region between I284 and V296 becomes disordered (Fig. 3a, Extended Data Fig. 10), suggesting that channel activation destabilizes this gating interface. Although the S4 and S5 helices in the Intermediate and Open states are shorter, this alone is not sufficient to cause pore opening. In the Open state, the S4 mid-helical bend is greater than in the Intermediate state. The dramatic secondary structure changes in the S4-S5 intracellular gating interface are not observed in the adjacent S6 helix, which retains its structure while being displaced radially. Taken together, these findings suggest that the combined unwinding and bending of the HCN S4 helix drive the conformational changes that ultimately lead to channel opening.

**Fig. 3:**
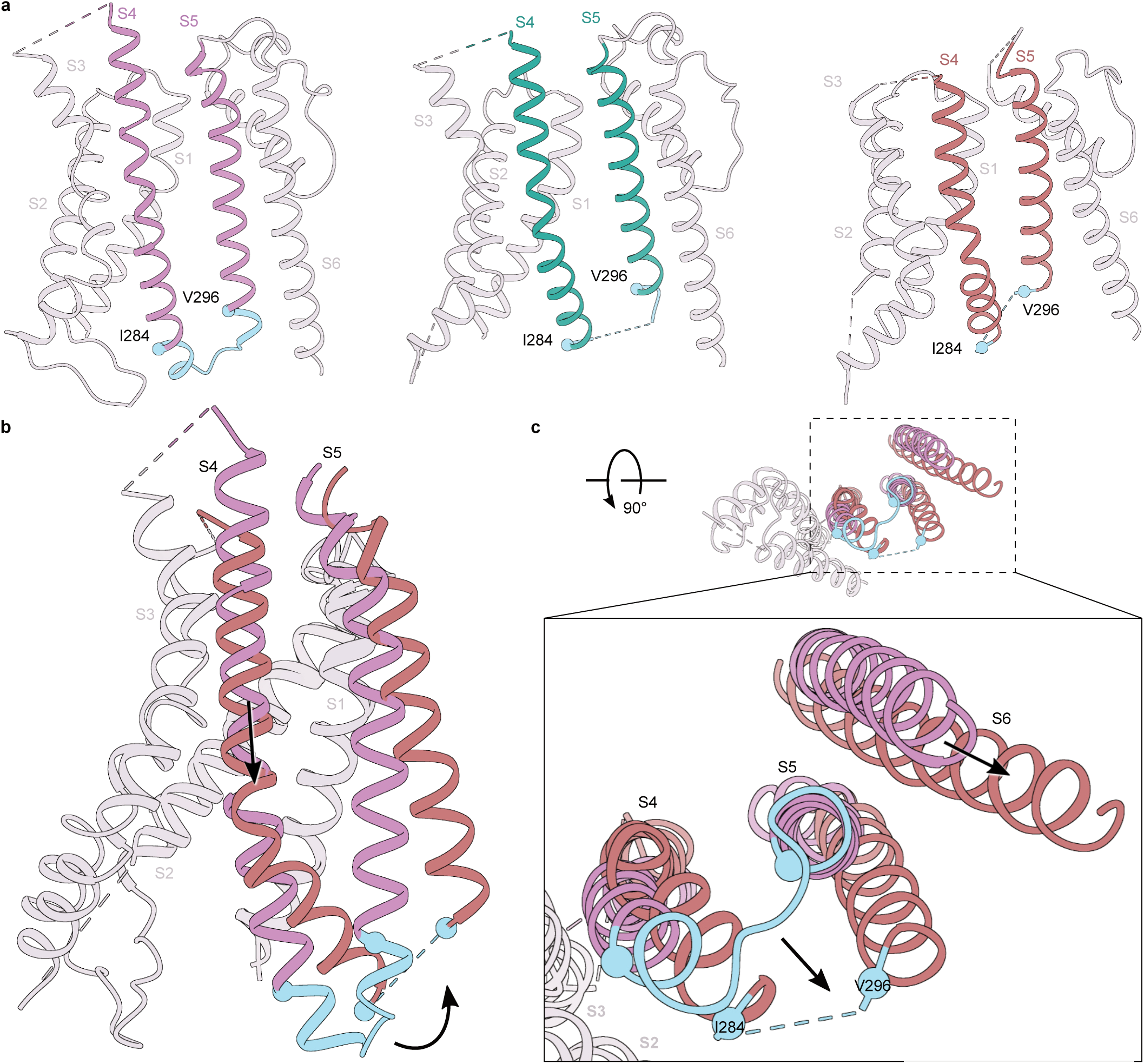
Conformational changes in the EM coupling interface. **a**, Structures of a protomer from Closed, Open and Intermediate states with S4 and S5 helices highlighted. The region between residues I284 and V296 is colored cyan. **b**, Comparison of the sideview of S1-S5 between the Closed and Open states. The P-loop and the S6 transmembrane helices were not depicted for clarity. **c**, Bottom view of transmembrane helices S1-S6 of Closed and Open states. Inset highlights the rearrangements of S4-S6 helices between the two states. Structures were aligned to S2.

While the central pore of the HCN channel is observed to be nonconductive in the Closed and Intermediate state, it is definitively dilated in the Open state (Fig. 4). Using the HOLE software, we were able to obtain a quantitative estimate of the width of the ion conducting pore. Fig. 4 shows that the Closed structure has two major constrictions: one in the selectivity filter and the other in the bottom intracellular gate. In both the Closed and Intermediate structures, the diameter of the pore gate is too small to allow passage of any hydrated ions. The Intermediate state structure showed a marginal increase in the diameter of the pore, but without any noticeable rotation of the S6 transmembrane helices. However, the diameter of the pore gate in the Open state structure is about 5.8 Å, in contrast to only 1.1 Å of that in the Closed state structure. The outward displacement of the S6 helix and rotation of the gate residues Y386, V390, and Q398 away from the pore axis facilitate pore widening. It is important to note that the rearrangement of S6 requires concerted movement of S5 helices to prevent steric clashes. Although the pore diameter in the selectivity filter does not change significantly, it does undergo rearrangement concomitant with gating transitions at the pore gates. Notably, the P-helices rotate clockwise and shift downwards to fill the growing gap between the upper portion of adjacent S6-helices upon channel opening. It is worth noting that the HCN channel pore is only weakly selective to potassium than sodium ions, and it functions as a non-selective monovalent cation-conducting pore.

**Fig. 4:**
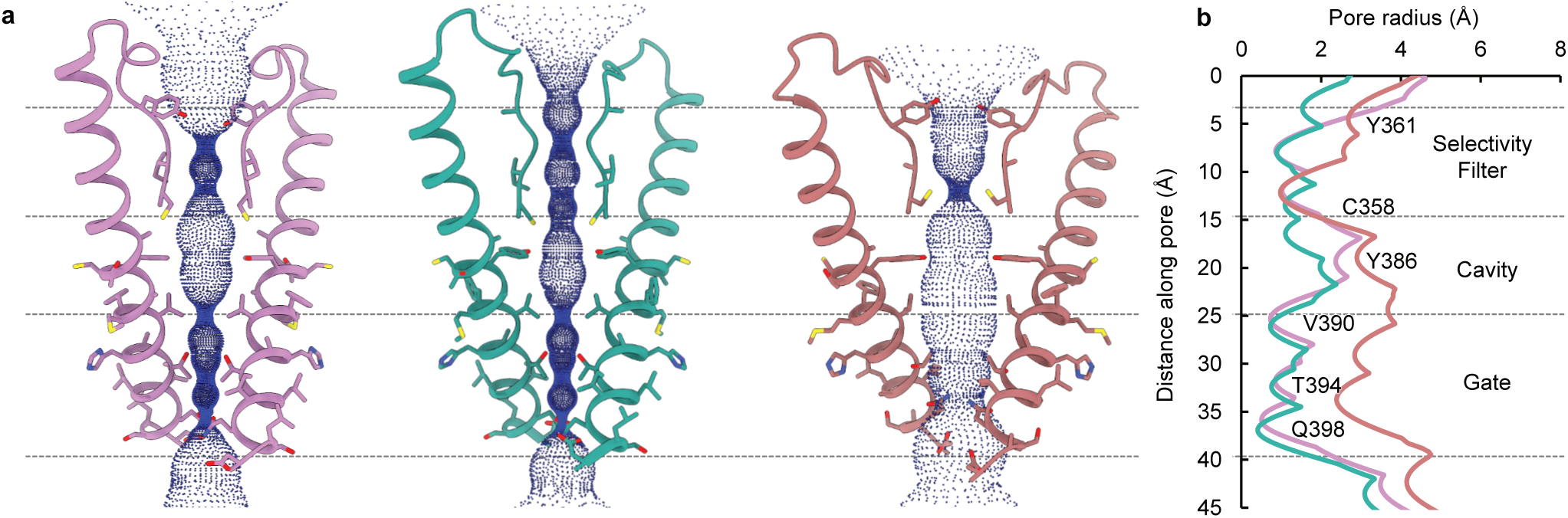
Ion conduction pathway. **a**, Comparison of the solvent accessible pathways in the Closed, Intermediate and Open states. The pore lining S6 helices and the selectivity filter of two opposing subunits (ribbon) are depicted together with the corresponding solvent accessible pathway generated by the HOLE program. Pore-lining residues are shown as sticks. **b**, Plot of the pore radii for all three structures.

All inwardly rectifying ion channels in the VGIC superfamily comprise a conserved CNBD that follows the six-transmembrane helices on the C-terminal end. While in HCN channels, the binding of cyclic nucleotides to this domain promotes voltage-dependent opening, the role of this domain in other hyperpolarization-activated ion channels such as KAT1 channels remains unclear^27^. The effect of cAMP binding on voltage-dependent activation is isoform-dependent with a voltage-sensitivity shift of ∼15-30 mV in HCN2 and HCN4 and an ∼5 mV shift in HCN1^28–31^. To study the impact of voltage-dependent gating on structural transitions in the CNB domain, all three structures were also obtained in the presence of a saturating concentration of cAMP to eliminate the confounding effects of ligand activation. The cryo-EM density maps of the cAMP-bound CNBD are in general of higher resolution than the transmembrane regions and are similar to those described in previous cryo-EM structures of HCN channels^17,19,21,32^. The structures of the CNBDs in the Closed, Intermediate and Open states are nearly identical (Fig. 5). The four cyclic nucleotide binding domains assemble into a symmetric gating ring below the pore, which is stabilized by extensive inter- and intra-subunit interactions in the C-linker region that connects the CNBD to the S6 pore helix (Fig. 5a). Pore opening causes a slight expansion of the CTD mainly due to the rearrangement of the C-linker region (Movie 4). Local superposition of the isolated C-linker and CNBD reveals a few notable structural changes in the CNBD upon pore opening (Fig. 5b). The inter- and intra-subunit interface of the C-linker region and CNBD region changes upon channel gating. The primary difference between the Intermediate and Open state is further displacement of the A’ helix in the C-linker region. The C-terminal end of the E’ helix interacts with one of the strands on the β-jelly roll in the Closed conformation, but in the Open state structure, this helix is poorly resolved, and there is a loss of density in the interacting strand.

**Fig. 5:**
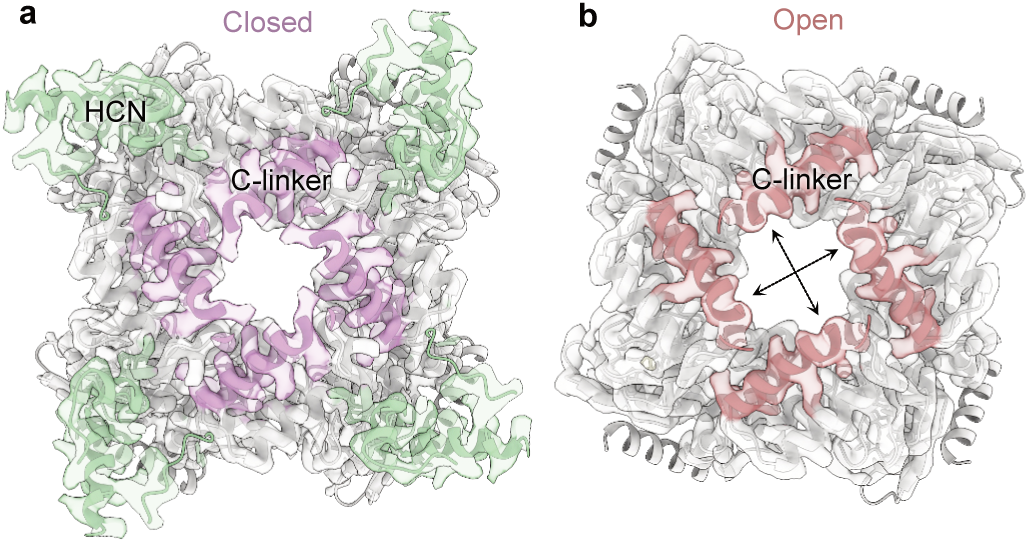
CTD ring during channel gating. **a-b**, Top view of the intracellular domain containing the C-linker (A’-F’ helices) and CNBD (A-E helices and β-roll) in the Closed (**a**) or Open (**b**) conformations, respectively. Protein structures and corresponding density maps are depicted. The distal E-helix is resolved only in the closed conformation. b, Comparison of intracellular CTD domain (sideview). HCN domain was only resolved in the Closed state and is colored green. The structures were globally aligned and the A’-helix of the C-linker are colored according to the same color scheme used previously. Arrows indicate expansion of the CTD ring.

## DISCUSSION

Most of the structures of hyperpolarization-activated ion channels published to date correspond to channels in a closed state where the voltage sensor is in the resting (Up) conformation^17,21,25^. These structures share an unusual feature that hints at the mechanism of reversed gating polarity in these ion channels. Specifically, the S4 helix in these structures is notably longer than S4 segments in depolarization-activated ion channels and protrudes to the intracellular side, where it interacts with the arm of the C-linker from a neighboring subunit. This characteristic feature has been observed in both plant KAT1 and animal hyperpolarization-activated (HCN) ion channels. It has led to the hypothesis that the interactions between the lower end of the long S4 helix and the C-linker region stabilize the closed pore and are critical for electromechanical coupling. Thus, the C-linker serves as a key mediator of electromechanical coupling analogous to the S4-S5 linker helix in the domain-swapped ion channels^33,34^. However, it is worth noting that truncated HCN channels, where the entire C-terminus is deleted, including the C-linker and CNBD, exhibit wild-type voltage-dependent activity and show no apparent loss of electromechanical coupling^29^.

Our structures of the human HCN1 isoform uncover several features that are common in both Open and Intermediate conformations. Firstly, the S4 helix is bent at the middle but not as much as in the activated voltage-sensor pre-open structure of HCN channels (PDB: 5U6P)^19^. In fact, the bend angle is more comparable to that of the S4 helix in other channels like TAX-4 and EAG1, both of which belong to the CNBD family (Extended Data Table 2)^18,35^. The bend angle is also remarkably similar to that of the predicted activated state structure based on FRET measurements^36^. Second, the S4 and S5 helices in both structures have lost secondary structure at the intracellular end, resulting in increased physical separation between the helices and disruption of the tight intracellular gating interface. Mutations near this interface increase voltage-dependent open probability, indicating destabilization of the closed state. For instance, a salt bridge formed by residues R297 and D401, which is important for stabilizing the closed state, is abrogated upon activation^34,37^. Third, the C-linker does not interact with the S4-S5 loop in the Open or Intermediate state but does so in the Closed state. Finally, voltage-dependent pore opening is accompanied by rearrangements in tetramerization interface mediated by the C-linker and CNBD. These findings are consistent with functional data that suggest that although the C-terminal domain indirectly facilities pore opening, it is not critical for gating polarity^29^.

Another surprising finding is the role of interaction between S5 and S6 in channel opening. Specifically, the C309-C385 disulfide bond formation is only observed in the Open state, but not in the Closed state. Preventing the formation of this bond by mutating C309 causes the pore to close, supporting the notion that – at least structurally – this bond is essential to stabilizing the Open state. Our functional data also suggests that the disulfide bond may be important for stability of the open pore. The disulfide bond in our structure is highly strained which is a hallmark of allosteric disulfide bonds^38,39^. The calculated disulfide strain energy of C309-C385 is 34.7 kJ/mol in comparison to 10 kJ/mol for a typical structural disulfide bond (Disulfide Bond energy server, https://services.mbi.ucla.edu/disulfide). Such high-strain disulfide bonds have been found to regulate the activity of a variety of enzymes, are highly labile, and their formation is conformation-specific^40^.

By comparing the structures of HCN channels in the three different conformations, we propose a mechanism of reversed electromechanical coupling. An applied voltage moves the voltage sensor downward past the charge transfer center, causing the intracellular ends of the S4 and S5 helices to unwind in a concerted manner, leading to the disruption of the tight intracellular gating interface (Movie 5). The loss of interactions at this interface allows the bottom half of the S5 helix to separate, which creates a space for the S6 helix to move into, resulting in pore dilation (Movies 5 and 6). The diameter of the gate residue Q398 increases to approximately 6 Å which is sufficient for hydrated potassium ions to permeate through the pore, as demonstrated in HCN4 channel through MD simulation^21^. Surrounding lipid molecules are likely to play a major role in stabilizing the intracellular gating interface^21,25,41^. We can resolve lipids in the Closed state structure but not in the activated conformation. It is unclear whether these differences are simply due to lower resolution of Open and Intermediate state structures or reflects state-dependent displacement of interacting lipids. Sea urchin HCN channels (spHCN) can be trapped in closed and open conformations by cross-linking of the S4-S5 linker with C-linker region^42–45^. The limited direct interaction between the C-linker and S4-S5 linker observed in our structures suggests that these crosslinks in HCN channels may simply reposition the C-linker, resulting in stabilization of the preceding S6 pore helix in one or the other conformation depending on the free-energy landscape.

To gain insight into mechanisms that result in opposing gating polarities, we compared the recently reported resting state structures of EAG1 (PDB: 8EP1)^18^ channel with our structure of the activated, open HCN channel. The most notable difference between the two channel types is how the S4-S5 helices interact with each other in a state-dependent fashion. In EAG channels, the S4 helix is of standard length and in Down conformation, the S4 helix is positioned close to the S5 due to interaction between the two helices and keeps the pore closed at negative voltages. In HCN channels, on the other hand, the extended S4 helix forms a tight interface with the S5 and stabilizes the closed pore conformation at depolarized potentials. When the membrane is hyperpolarized, the downward movement of HCN S4 disrupts the intracellular gating interface largely due to a loss of secondary structure in the lower part of the S4-S5 helices. This loss of stabilizing interactions initiates further rearrangements of the S5 and S6 helices culminating in pore opening.

While the concerted unwinding of multiple helices during channel gating is unusual, individual transmembrane helices have been shown to undergo helix-coil transitions. For example, the S6 helix of TRPM8 channels undergoes this transition during activation^46^. Similarly, the intracellular end of S4 helix of Nav1.7 channel also unwinds when the voltage sensors move to the Down conformation^24^. Additionally, the intracellular ends of the HCN S4 and S5 helices are situated outside the bilayer, particularly at hyperpolarized potentials, making this secondary structure transition energetically less unfavorable. This transition is less expensive in aqueous medium than in the membrane, where there are no water molecules to form hydrogen bonds with the backbone amides. It is also possible that some of the energetic costs associated with these structural transitions result in weak allosteric coupling between the voltage sensor and pore in these channels. Nevertheless, it is clear that further experimental evidence is necessary to evaluate the proposed model of HCN channel activation and determine if this mechanism applies to other hyperpolarization-activated ion channels.

## MATERIALS AND METHODS

### Molecular Biology and Protein purification

The human HCN1-EM construct in the pEG vector was used as a background for all the mutants in this study^17^. The C-terminal region spanning amino acids 636-865 are deleted and the N-terminus is tagged with eGFP followed by a HRV 3C protease cleavage site. We added a twin-strep tag on the N-terminus for affinity purification^47^. HCN1-EM-F186C-S264C and HCN1-EM-F186C-S264C-C309A mutants were made using the FastCloning technique^48^ and verified by Sanger sequencing before use.

Both HCN1-EM-F186C-S264C or HCN1-EM-F186C-S264C-C309A were expressed in suspension cultures of Freestyle HEK cells using a BacMam system^49^. Bacmid DNA containing these constructs were amplified in DH10Bac competent cells (Bac-to-Bac; Invitrogen). Sf9 cells were transfected with Bacmid DNA (1-2 µg/10^6^ cells) using Cellfectin II (Invitrogen). The supernatant containing P1 baculoviruses was harvested after 5 to 7 days, sterile filtered and used to generate P2 baculovirus (dilution factor: 100). The P2 baculovirus was sterile filtered and supplemented with 2.5 % FBS before use. For transduction, 800 mL suspension culture of Freestyle HEK cells (3-3.5×10^6^ cells/mL) were transduced with 3.5-5% baculovirus. After 10-12h at 37 °C, the cells were supplemented with 10 mM sodium butyrate and kept at 30 °C for another 48-50 hours. The cells were harvested by low-speed centrifugation and cell pellets were stored at -80 °C.

For purification, the cell pellet of 400-1200 mL suspension culture was lysed with 50-80 mL hypotonic lysis buffer (20mM KCl, 10mM Tris, protease inhibitor cocktail (Sigma Aldrich, P8340), pH 8.0) and sonicated 6-8 times for 10 s. The membrane fraction was collected by spinning at 50,000 g for 45 mins. The membrane pellets were solubilized in 70 mL solubilization buffer (300 mM KCl, 40 mM Tris, 4 mM DTT, 10 mM LMNG, 4 mM CHS, protease inhibitor cocktail (P8340, Sigma Aldrich), pH 8.0) for 1.5 h. After removing the insoluble fraction by centrifugation, the supernatant was incubated with Strep-Tactin Sepharose (iba, #2-1201-025) for 2.5 hrs, packed onto a column and then washed with 3.5 column volumes of wash buffer. The protein was eluted in 3-6 mL wash buffer containing 10 mM d-desthiobiotin. After removing Twin-Strep-tag and GFP using 3C-protease, the protein was further purified using a Superose 6 Increase 10/300 GL column (Cytiva Life Sciences, # 29091596). . For obtaining the closed state structure, the SEC buffer was supplemented with 2 mM DTT. For all samples, 200 µM HgCl was added, and the protein samples were concentrated to 1-2.5 mg/mL using a concentrator (cut-off: 100 kDa, Amicon Ultra, Millipore). Representative SEC profiles and Coomassie blue stained SDS-PAGE gels for the Closed, Intermediate and Open state structure are depicted in Extended Data Fig. 11.

### Cryo-EM grid preparation and image acquisition

Quantifoil R1.2/1.3 or R2/2 holey carbon copper grids were cleaned for 60 s using a Gatan Solarus 950. Before plunge freezing, the samples were spiked with 1 mM fluorinated fos-choline (Anatrace, F300F) and incubated for 1 min at 100% humidity and 4° C using a FEI Vitrobot Mark IV.

The closed-state structure was obtained from a single dataset collected on a 200 kV Glacios TEM equipped with a Falcon IV detector (ThermoFisher Scientific). Movies were collected at a magnification of 150,000 X. The calculated pixel size was 0.94 Å and the nominal defocus was set between -0.8 and - 2.4 µm. A dose rate of approx. 5.43 e^-^/Å^2^/s was used with a cumulative dose of 53.07 electrons per Å^2^. The intermediate and the open state structures were obtained from four datasets collected on a 300 kV Titan Krios TEM equipped with a Falcon IV detector at a magnification of 59,000 X. The calculated pixel size was 1.16 Å and the nominal defocus was set between -0.8 and -2.4 µm. The dose rate used was in the range of 3.99-4.74 e^-^/Å^2^/s and did not exceed a cumulative dose of 55.07 electrons per Å^2^.

### Cryo-electron microscopy image processing

A detailed overview of the data-processing workflow of the Closed, Intermediate and the Open state structure is provided in Extended Data Figs. 1, 4 and 8. In all three cases, movie frames were aligned and dose weighted using patch-based motion correction, and then subjected to patch-based contrast transfer function (CTF) determination in CryoSPARC 4.1^50^. After motion correction^51^ and CTF estimation^52^, manual curation of the dataset was performed on the basis of estimated CTF resolution, cross-correlation, and ice contamination. Particles were picked using a blob-based autopicker within a 150Å to 300Å range and subjected to 2D classification. 2D classes corresponding to intact HCN1 channels were combined and used to generate an *ab initio* model with C1 symmetry, which was subsequently refined with C4 symmetry, lowpass filtered to 15 Å, and used as a template for heterogeneous refinement. These classes underwent non-uniform refinement^53^, revealing either junk classes or intact channels. The classes corresponding to intact channels were combined, refined together with C4 symmetry and utilized to generate 2D projections corresponding to all possible channel orientations. These projections were lowpass filtered to 20Å to prevent Einstein from noise or model bias and were used for particle re-picking resulting in slightly higher particle count and improved orientational sampling. If no prominent lipid density was present in the final map, DeepEMhancer^54^ was utilized to sharpen the final half-maps.

For the Closed conformation, 1,973 exposures were manually curated to remove low-quality micrographs, resulting in 1,934 that were retained for further processing. Blob-based picking identified 610,860 particles which were subjected to 2D-classification to identify particles representing intact channels. These 93,472 particles were utilized to generate templates for particle re-picking. Template-based picking identified 698,204 particles which were subjected to 2D classification – 103,753 were retained for further processing. Particles were subsequently separated into three classes using heterogeneous refinement with C4 symmetry. Two of the classes represented intact HCN1 channels and were essentially identical, while the remaining class had broken density, was at much lower resolution, and was discarded. The two identical classes were combined, subjected to nonuniform refinement with C4 symmetry imposed, and underwent iterative per-particle CTF-refinement defocus correction until map quality could no longer improved. The final 88,923 particles were refined to 3.26 Å resolution as assessed by gold-standard Fourier shell correlation (FSC).

To obtain the Intermediate conformation of HCN1, 2,248 of 2,575 exposures were used for downstream analysis. Blob-based particle picking as described above identified 905,636 particles, 168,944 of which were retained after 2D-classification. 108,901 of these particles were used to generate templates for particle re-picking. Template-based picking identified 859,495 particles, 188,999 of which were kept for downstream analysis. After heterogeneous refinement with C4 symmetry with 4 classes, 3 had broken density and were at much lower resolutions than the fourth class which was retained for further processing. The final 77,983 particles were refined with nonuniform refinement to a 3.89 Å resolution as assessed by gold-standard FSC. CTF-refinement, defocus correction, and local motion correction did not improve the quality of the map. The final half-maps were sharpened using DeepEMhancer on the high-resolution preset.

For the HCN1 open conformation, 8,196 movies were collected, 7,828 of which were utilized. Blob picking was performed as described above, and 710,625 particles corresponding to intact channels were identified after subsequent template-based particle repicking. Particles were separated into six classes using heterogeneous refinement. 3 classes corresponded to channels with severely broken density, Two classes, consisting of 288,025 particles, corresponded to channels with intact features were combined. To improve the resolution in the TM-helices, a mask surrounding the transmembrane region and channel core was manually generated for focused classification. The particles and the manually generated mask were imported into Relion 4.0^55^ and subjected to 3D-classification without orientational assignment to identify channels with improved TM density. After separation into 4 classes, 2 classes with the highest resolution in the S6 helix were combined, and the corresponding 255,148 particles were imported into CryoSPARC for nonuniform refinement with C4 symmetry, resulting in a 3.73 Å map. The final half-maps were sharpened using DeepEMhancer on the high resolution preset.

### Model building and validation

For building the initial model, the cAMP-bound HCN1 closed state structure (PDB: 5U6P)^19^ was rigid-body fitted into the cryo-EM density map in a segment-based manner using COOT^56,57^. Unresolved regions were deleted from the model, and secondary structure features and side chains not fitting the density were manually rebuilt and refined in COOT. To avoid overfitting, residues with missing side chain density were trimmed. In iterative cycles, the atomic models were manually corrected for side-chain outliers in COOT and real-space refined in PHENIX^58^. The final atomic models were validated using MolProbity^59^ and the RCSB PDB validation server. Cryo-EM data collection and refinement statistics are summarized in Extended Data Table 1. Structural illustrations were prepared with UCSF ChimeraX^60^.

### Heterologous Expression and electrophysiology

hHCN1-EM or hHCN1-F186C-S264C-C309A were subcloned into a pUNIV vector^61^ and cRNAs were synthesized using a mMessage T7 transcription kit (Invitrogen). Isolated Xenopus laevis oocytes were injected with 20-40 ng cRNA and incubated at 17° C for 24-48h before recording. Two-electrode voltage-clamp (TEVC) recordings were obtained at room temperature using a CA-1B amplifier (Dagan) at a sampling rate of 10 kHz and were filtered with a cutoff frequency of 5 kHz. Electrodes were fabricated from thin-walled glass pipettes (World Precision Instruments) using a P97 micropipette puller (Sutter Instruments). The pipette resistance was in the range of 0.5-0.9 MΩ using 3 M KCl. The bath solution contained 107 mM NaCl, 5 mM KCl, 20 mM Hepes, and 2 mM MgCl_2_ (pH = 7.4 NaOH). For channel activation, the oocytes were held at a holding potential of -20 mV and were then conditioned to potentials ranging from -20 mV to -110 mV with 10 mV decrements before pulsed to 0 mV at which the tail currents were recorded. The leak current-corrected peak amplitude of the tail current reflects the fraction of activated channels and was plotted as a function of the conditioning pulse. The activation curve was normalized and fitted with a single Boltzmann function:

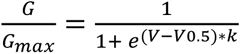

where V is the conditioning pulse, V_0.5_ is the potential of half maximal activation, k is the slope factor, a is the peak amplitude of the tail current and o is the offset current. Data were collected and analyzed using pCLAMP10.0 (Molecular Devices) and plotted using OriginPro 2020b (OriginLab Corp., Northampton MA).

Data shown in Extended Data Fig. 7 represent mean ± SEM. Mean half-maximal activation were compared between HCN1-EM, HCN1-EM-F186C-S264C and HCN1-EM-F186C-S264C-C309A using a nonparametric Mann-Whitney post-hoc test with a significance level of p < 0.05.

### Data availability

The cryo-EM density maps and the corresponding atomic models of the different gating conformations of the human HCN1-EM-F186C-S264C and the human HCN1-EM-F186C-S264C-C309A have been deposited to the Electron Microscopy Data Bank (EMDB) and the Worldwide Protein Data Bank (wwPDB). The accession codes for the closed-state conformation of hHCN1-EM-F186C-S264C are EMD-41036 and 8T4M; for the open-state conformation of hHCN1-EM-F186C-S264C are EMD-41041 and 8T50; and for the intermediate-state conformation of hHCN1-EM-F186C-S264C-C309A are EMD-41040 and 8T4Y, respectively.

## ACKNOWLEDGEMENTS

The research was supported by NIH grants to B.C. (NS101723) and P.Y. (GM143440). V.B. is supported by Schrödinger fellowship (J 4652) by Austrian Science Foundation. We thank R. MacKinnon for generous gift of hHCN1-EM pEG plasmid and C. Czajkowski for the pUNIV vector. We thank Drs. B.T. Summers, K. Basore and M. Rau at Washington University Center for Cellular Imaging (WUCCI) or grid freezing and cryo-EM data collection.

## SUPPLEMENTARY FIGURE LEGENDS

**Extended Data Fig. 1:**
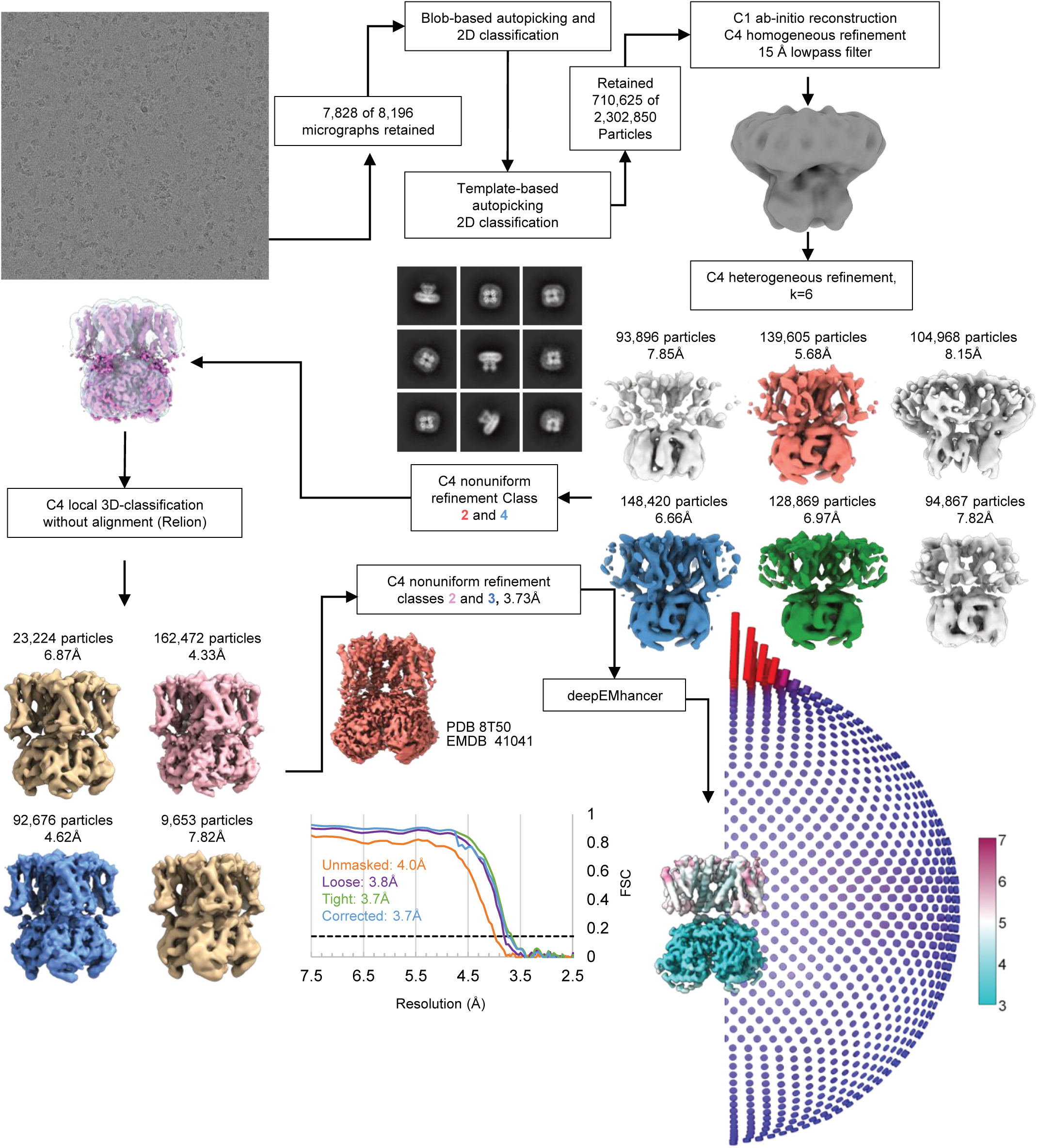
Cryo-EM processing workflow for the cAMP bound HCN1 Open conformation (HCN1-EM F186C-264C).

**Extended Data Fig. 2:**
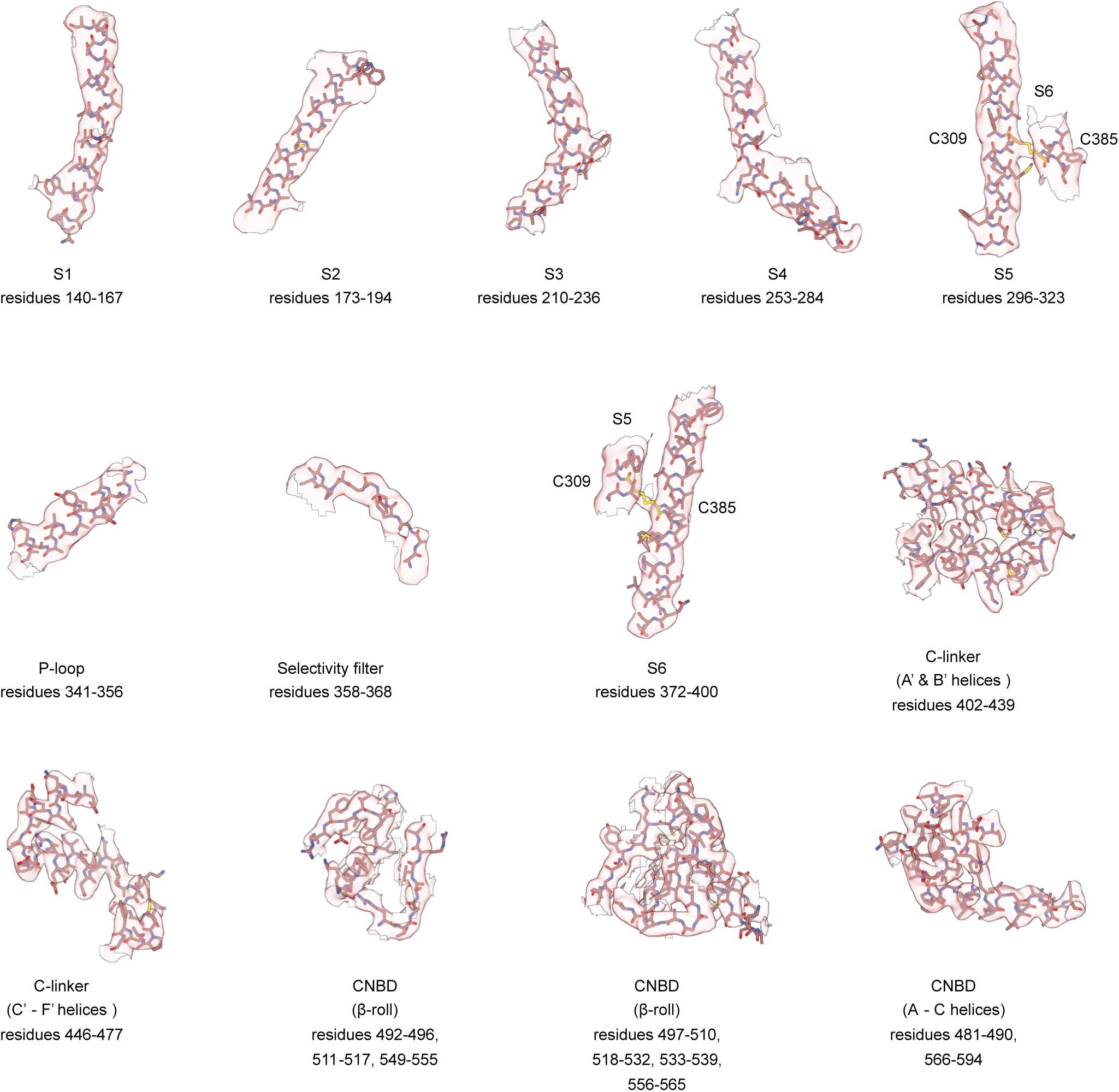
Cryo-EM density of cAMP-bound Open-state structure of HCN1-EM F186C-S264C. Structure and cryo-EM density maps are depicted for regions indicated.

**Extended Data Fig. 3:**
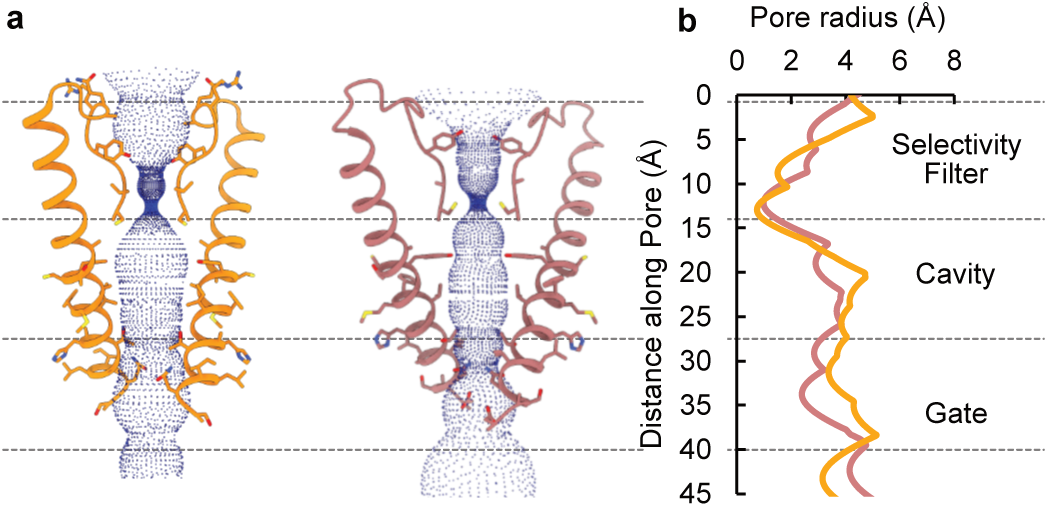
Ion conduction pathways of open HCN1 and HCN4. **a**, Comparison of the solvent accessible pathways of the open states of human HCN1 and rabbit HCN4. The pore lining S6 helices and the selectivity filter of two opposing subunits (ribbon) are depicted together with the corresponding solvent accessible pathway (blue dots) generated by the HOLE program. Residues comprising the pore are shown as sticks. **b**, Plot of the pore radii for both structures.

**Extended Data Fig. 4:**
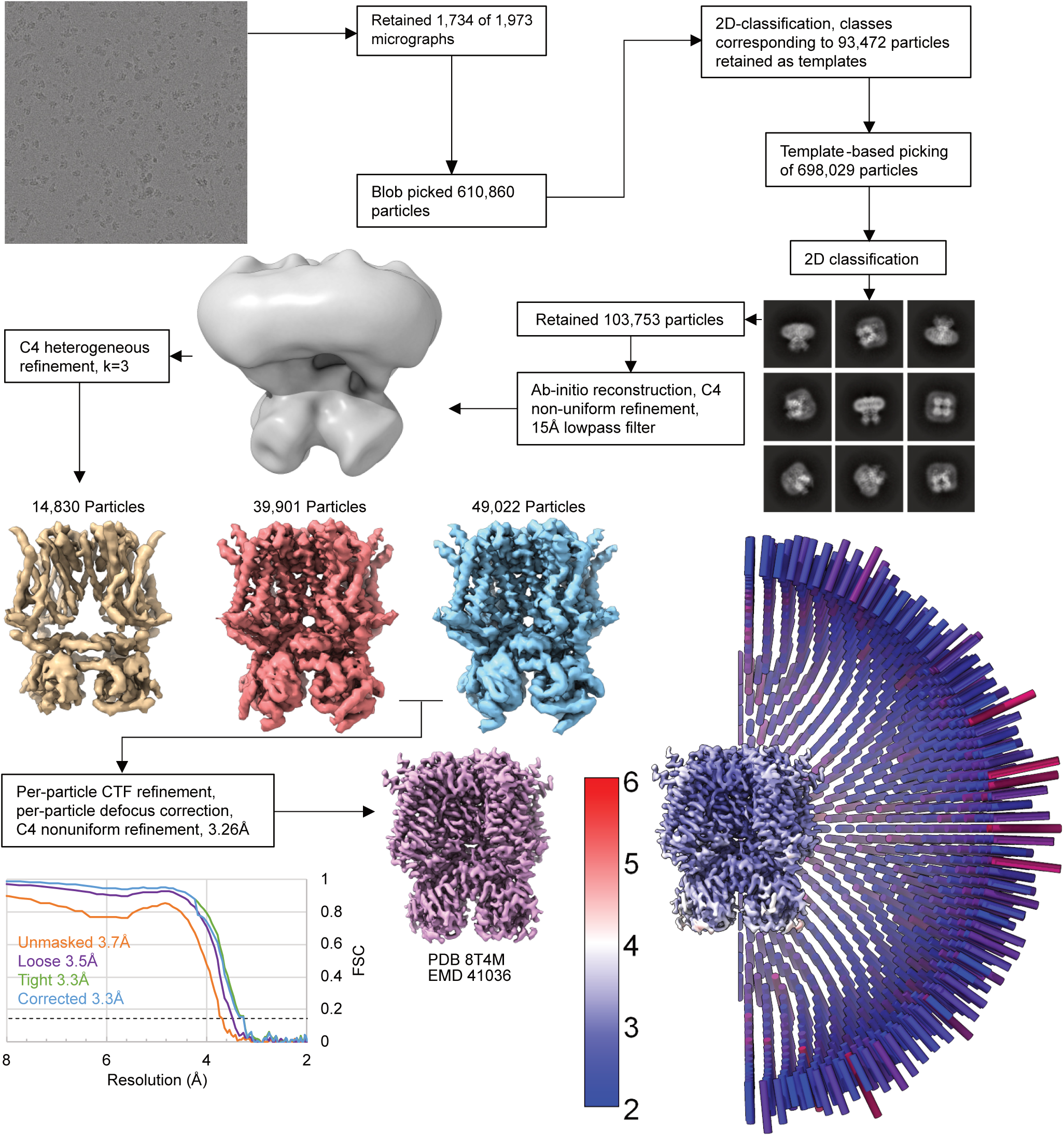
Cryo-EM processing workflow for the cAMP bound HCN1 Closed conformation (HCN1-EM F186C-264C).

**Extended Data Fig. 5:**
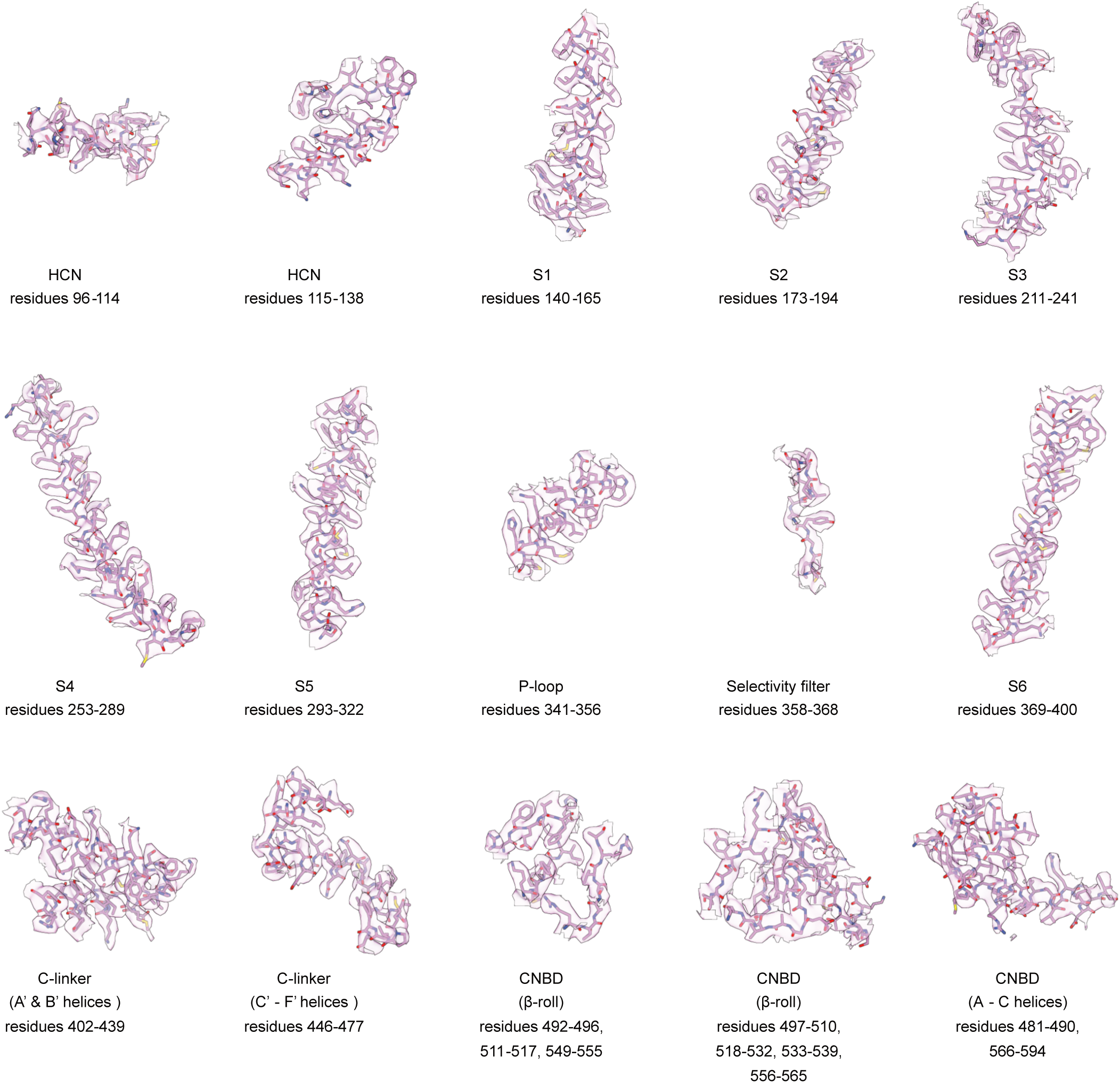
Cryo-EM density of cAMP-bound Closed-state structure of HCN1-EM F186C-S264C. Structure and cryo-EM density maps are depicted for regions indicated.

**Extended Data Fig. 6:**
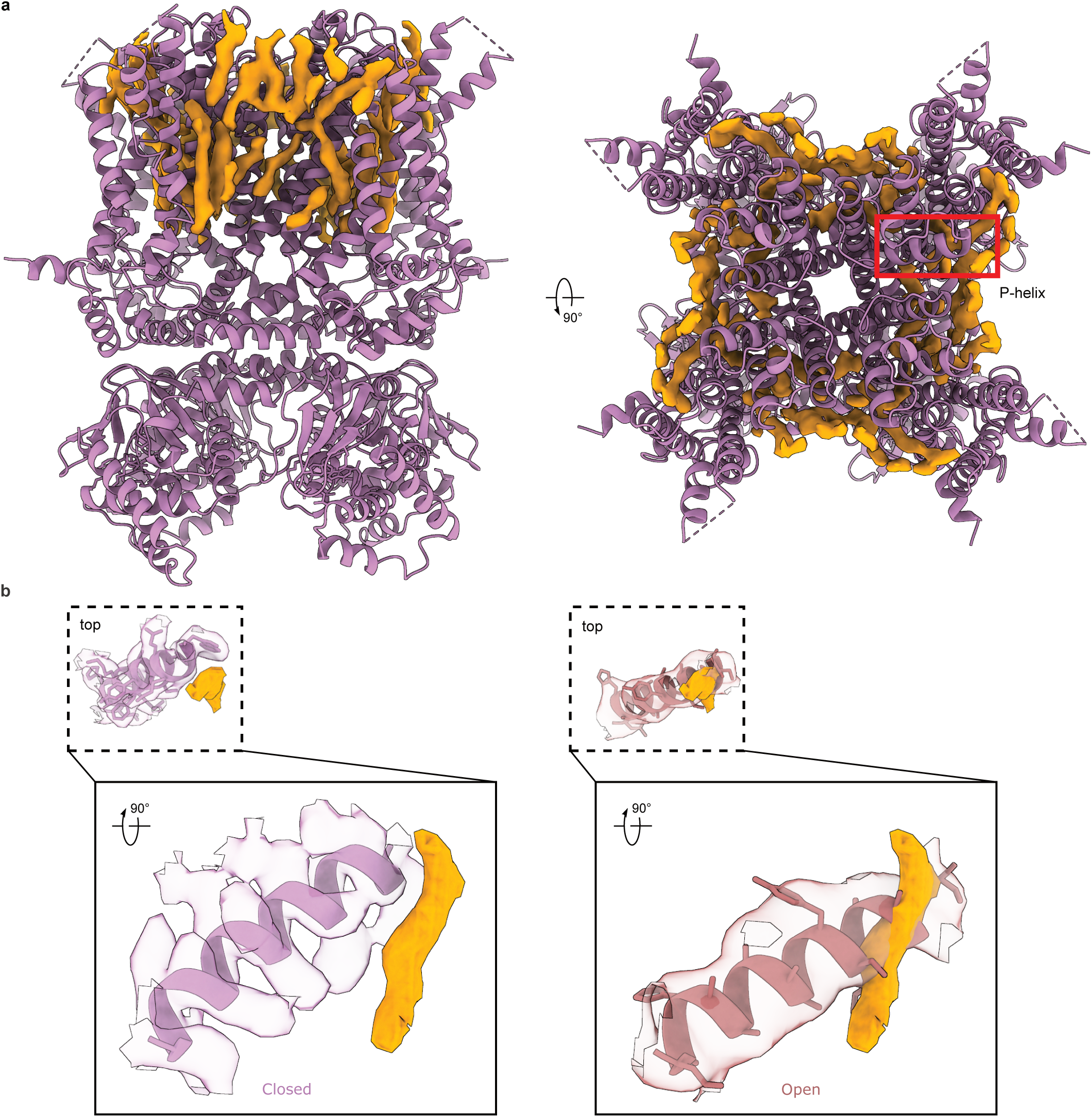
Annular lipids in the Closed state structure. **a,** The lipid density (highlighted in orange) in the Closed structure (lilac). **b**, Superposition of the lipid density observed in the Closed state on to the Open structure (burnt red) indicates overlap with the P-helix, where the lipids observed in the Closed structure would sterically clash with the Open channel. Inset in panel **b** depicts top view of the P-helix and neighboring lipid density.

**Extended Data Fig. 7:**
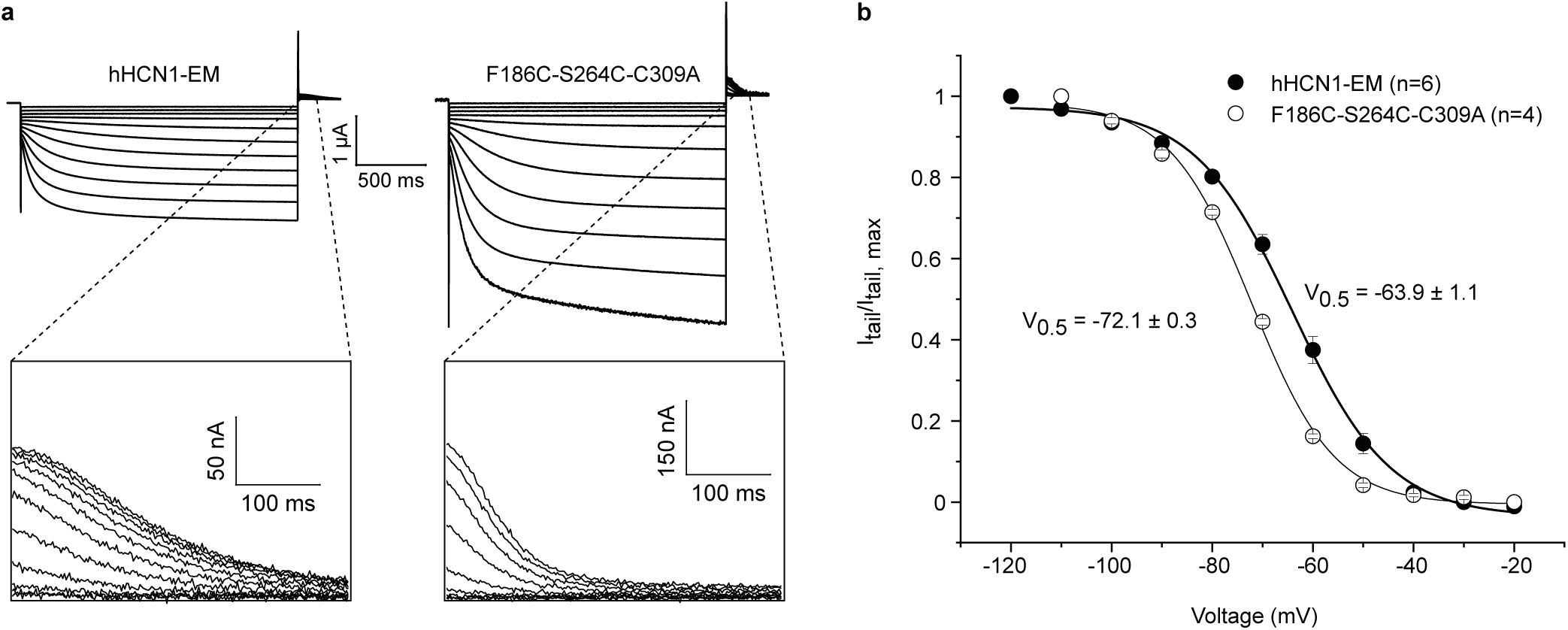
*h*HCN1-EM-F186C-S264C-C309A channels activate at more negative potentials and have faster deactivation kinetics than *h*HCN1-EM. **a**, Representative sample traces of HCN1-EM (left panel) and F186C-S264C-C309A (right panel) and corresponding close-ups of the tail currents. **b**, Conductance-voltage curves of F186C-S264C-C309A channels compared to HCN1-EM.

**Extended Data Fig. 8:**
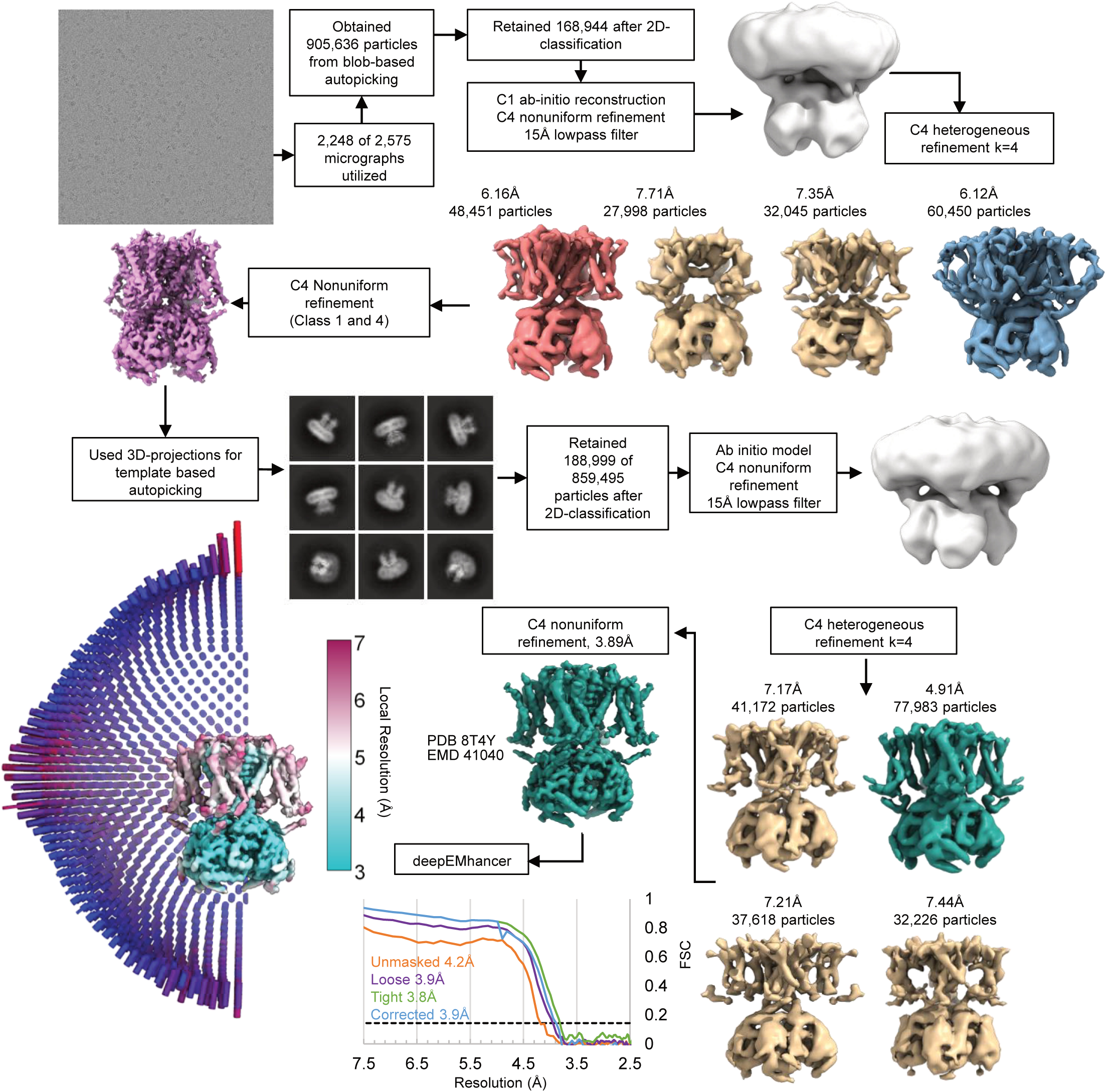
Cryo-EM processing workflow for the cAMP bound HCN1 Intermediate conformation (HCN1-EM F186C-S246C-C309A).

**Extended Data Fig. 9:**
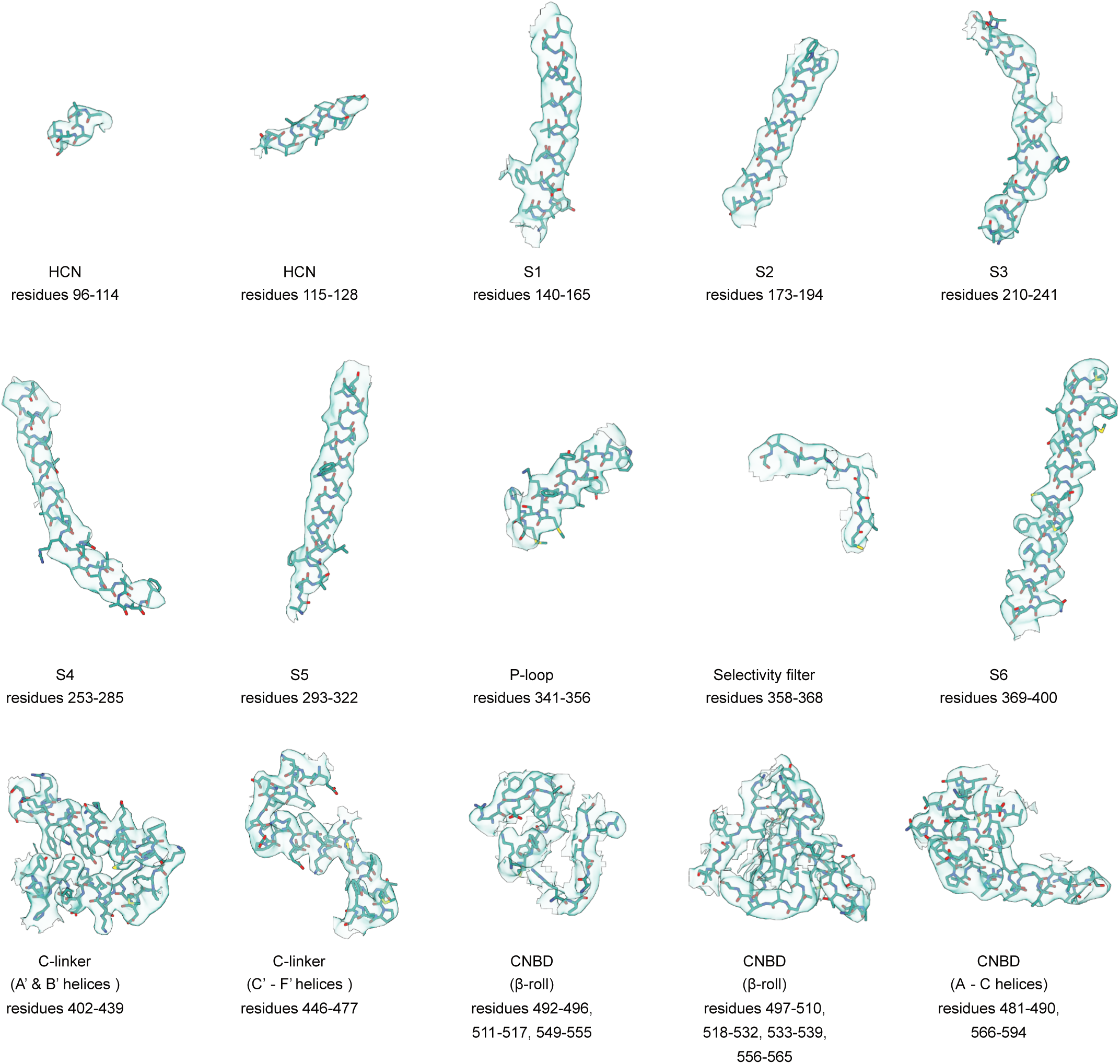
Cryo-EM density of cAMP-bound Intermediate-state structure of HCN1-EM F186C-S264C-C309A. Structure and cryo-EM density maps are depicted for regions indicated.

**Extended Data Fig. 10:**
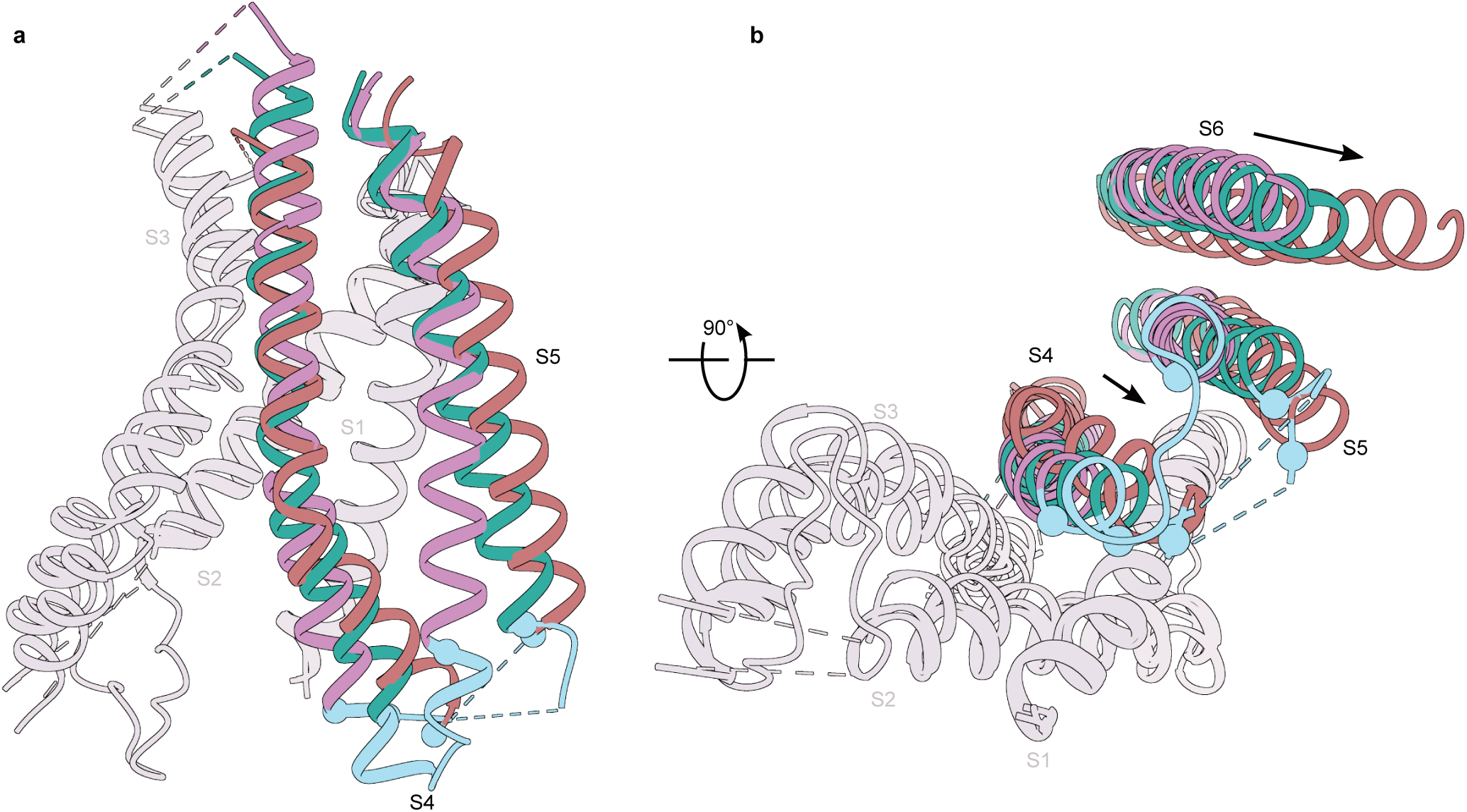
Conformational changes in the EM coupling interface of Closed, Intermediate and Open state. **a,** Sideview of transmembrane helices S1-S5. The region between I284 and V296 is shown in cyan and cannot be resolved in the Open state. The P-loop and the S6 transmembrane helices were not depicted for clarity. **b,** Bottom view of transmembrane helices S1-S6 of Closed, Intermediate and Open states. The S4-S6 helices undergo rearrangement between these states. Structures were aligned to S1 and S2 helices. The P-loop is not depicted.

**Extended Data Fig. 11:**
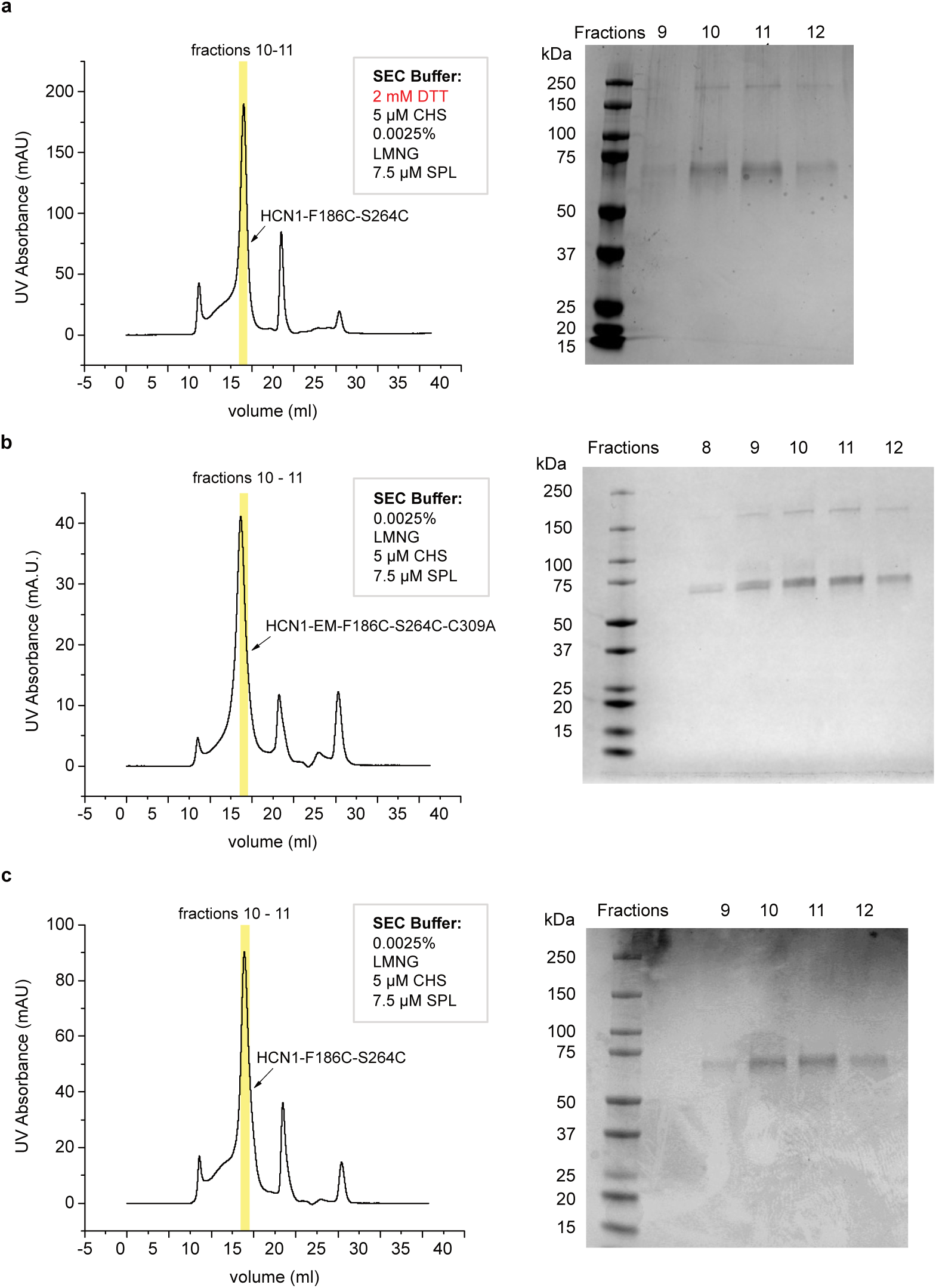
Representative size-exclusion chromatographs and Coomassie blue stained SDS-PAGE gels of the protein samples for obtaining the Closed, Intermediate and Open conformation. **a**, The Closed state was obtained from HCN1-EM F186C-S264C purified in LMNG/CHS/SPL in the presence of the reducing agent, dithiothreitol (DTT). **b**, The Intermediate state was obtained from HCN1-EM F186C-S264C-C309A purified in LMNG/SPL without DTT. **c**, The Open state was obtained from HCN1-EM F186C-S264C purified in LMNG/CHS/SPL without DTT. In all cases, fractions 10 & 11 (yellow) were pooled and used for grid freezing.

**Supplementary Table 1.** Cryo-EM data collection, refinement and validation statistics.

|  | *h*HCN1 Closed<br>EMDB-41036<br>PDB 8t4m | *h*HCN1 Intermediate<br>EMDB-41040<br>PDB 8t4y | *h*HCN1 Open<br>EMDB-41041<br>PDB 8t50 |
| --- | --- | --- | --- |
| **Data collection and processing** |  |  |  |
| Magnification | 150,000x | 59,000x | 59,000x |
| Voltage (kV) | 200 | 300 | 300 |
| Electron exposure (e-/Å<sup>2</sup>) | 53.08 | 55.07, 53.3 | 54.55, 53.99 |
| Defocus range (μm) | -0.8 to -2.4 | -0.8 to -2.4 μm | -0.8 to -2.4 μm |
| Pixel size (Å) | 0.94 | 1.1 | 1.1 |
| Symmetry imposed | C4 | C4 | C4 |
| Initial particle images (no.) | 698,029 | 859,495 | 2,302,850 |
| Final particle images (no.) | 88,923 | 77,983 | 255,148 |
| Map resolution (Å) | 3.26 | 3.89 | 3.73 |
| FSC threshold | 0.143 | 0.143 | 0.143 |
| Map resolution range (Å) | 2-4 | 3-6 | 3-6 |
| **Refinement** |  |  |  |
| Initial model used (PDB code) | 5u6p | 8t4m | 6uqf |
| Model resolution (Å) | 3.46 | 4.11 | 3.93 |
| FSC threshold | 0.5 | 0.5 | 0.5 |
| Model resolution range (Å) | 3.09-3.53 | 3.73-4.18 | 2.88-3.95 |
| Map sharpening *B* factor (Å<sup>2</sup>) | -158 | -184.7 | -223.9 |
| Model composition |  |  |  |
| Non-hydrogen atoms | 16952 | 13072 | 11864 |
| Protein residues | 2132 | 1936 | 1756 |
| Ligands | 4 | 4 | 4 |
| *B* factors (Å<sup>2</sup>) |  |  |  |
| Protein | 64.75 | 79.87 | 133.46 |
| Ligand | 110.13 | 62.97 | 110.33 |
| R.m.s. deviations |  |  |  |
| Bond lengths (Å) | 0.004 | 0.004 | 0.004 |
| Bond angles (°) | 0.942 | 0.996 | 1.014 |
| Validation |  |  |  |
| MolProbity score | 1.38 | 1.58 | 1.7 |
| Clashscore | 5.56 | 6.25 | 6.35 |
| Poor rotamers (%) | 0.28 | 0.55 | 0 |
| Ramachandran plot |  |  |  |
| Favored (%) | 97.64 | 96.45 | 94.96 |
| Allowed (%) | 2.36 | 3.55 | 5.04 |
| Disallowed (%) | 0 | 0 | 0 |

**Supplementary Table 2:** Conformation and bending angle of S4 helix in homologous CNG channels. Angles are calculated using anglebetweenhelices.py utilizing the Cα-fit preset.

| Channel | Conductive State | S4 conformation | Bending Angle | PDB |
| --- | --- | --- | --- | --- |
| *h*HCN1 | Closed | Up | 14.8° | 8t4m |
|  | Closed | Intermediate | 28.8° | 8t4y |
|  | Closed | Down | 60.7° | 6uqf |
|  | Open | Down | 35.9° | 8t50 |
| *r*HCN4 | Open | Up | 14.4° | 7np3 |
| *h*EAG | Closed | Down | 29.9° | 8ep1 |
| *c*TAX-4 | Closed | Down | 53.3° | 6wej |
| *c*TAX-4<sup>R421W</sup> | Open | Down | 54.9° | 7n17 |

## Notes

### Competing Interest Statement

The authors have declared no competing interest.

